# An mRNA–Lipid Nanoparticle Platform Encoding the Conserved Outer Membrane Protein BamA Elicits Broadly Cross-Reactive Systemic and Mucosal Antibodies Against Antimicrobial-Resistant *Neisseria gonorrhoeae*

**DOI:** 10.64898/2026.07.14.738439

**Authors:** Natalie Wolske, Namratha Turuvekere Vittala Murthy, Abhishek Chanda, Junaid Nazir, Ryszard A. Zielke, Fabian G. Martinez, Jeonghwan Kim, Ravi Kant, Gaurav Sahay, Aleksandra E. Sikora

**Affiliations:** Department of Pharmaceutical Sciences, College of Pharmacy, Oregon State University, Corvallis, Oregon, United States; Department of Pharmaceutical Sciences, College of Pharmacy, Oregon Health & Science University, Portland, Oregon, United States; Computational Drug & Vaccine Discovery Laboratory, Faculty of Applied Sciences & Biotechnology, Shoolini University, Solan, Himachal Pradesh-173229, India; Vaccine and Gene Therapy Institute, Oregon Health & Science University, Beaverton, Oregon, United States

**Keywords:** mRNA vaccine, *Neisseria gonorrhoeae*, BamA, LNPs, gonorrhea, mouse model, epitope prediction

## Abstract

*Neisseria gonorrhoeae* (*Ng*) is the causative agent of gonorrhea, and the global spread of antimicrobial-resistant strains makes vaccine development a public health priority. Although messenger RNA (mRNA) vaccines have transformed protection against viral diseases, the platform remains in its infancy against pathogenic bacteria. Here, we evaluated the immunogenicity and protective efficacy of an mRNA–lipid nanoparticle (LNP) vaccine encoding the highly conserved outer membrane antigen BamA in the female mouse model of lower genital tract infection. We delivered BamA mRNA-LNPs via intramuscular (IM) or intranasal (IN) routes, with or without CpG ODN 2395, and measured antigen-specific antibody responses in serum and vaginal lavage samples. Both routes elicited robust BamA-specific antibodies that recognized diverse *Ng* isolates, including ceftriaxone-resistant strains. However, neither route accelerated bacterial clearance nor reduced bioburden, nor did either generate serum bactericidal activity. These findings show that BamA mRNA-LNPs are immunogenic but, as formulated, are not protective, and they pave the way for modifications to the construct, adjuvant, and route. To our knowledge, this is the first evaluation of an mRNA vaccine against *Ng*, establishing the platform as an amenable approach for gonococcal antigen testing.

**Importance:** *Neisseria gonorrhoeae* is a high-priority public health threat, and no licensed gonococcal vaccine exists. mRNA-lipid nanoparticle vaccines have transformed antiviral immunization but remain largely untested against bacterial pathogens. This study is the first to evaluate an mRNA vaccine against *N. gonorrhoeae*, using the conserved, essential outer membrane antigen BamA. We show that BamA mRNA-LNPs delivered by intramuscular or intranasal routes elicit robust, broadly cross-reactive antibodies that recognize diverse isolates, including ceftriaxone-resistant strains, even though the tested formulations did not confer protection in mice. By establishing mRNA-LNPs as a viable platform for gonococcal antigen testing, this work lays the foundation for optimizing constructs, adjuvants, and routes for developing mRNA-based vaccines against gonorrhea and other antimicrobial-resistant bacteria.

## Introduction

Recent advances in mRNA technology and delivery systems have ushered in a new era of vaccine development and therapeutic interventions against infectious diseases, cancer, and monogenic disorders (1). mRNA platforms enable time-efficient vaccine design, scalable manufacturing, and a favorable safety profile because their components are non-infectious and not derived from a living pathogen (2, 3). The successful outcomes of COVID-19 mRNA vaccination and subsequent studies demonstrated that mRNA vaccines are effective against viral infections and cancer (4). The use of mRNA technology, however, remains largely unexplored against pathogenic bacteria; to date, 16 preclinical studies have been published, and three vaccines have entered clinical evaluation (5). One of the first reported mRNA vaccines encoding *Mycobacterium tuberculosis* lipoprotein MPT83 conferred modest but significant protection against bacterial challenge following intramuscular immunization (6). Similarly, a single intranasal dose of mRNA Hsp65 from *M. leprae* elicited protection against a virulent strain of *M. tuberculosis* (7). The tuberculosis LNP-mRNA vaccines induced a Th1-polarized response and enhanced long-term protection when used with BCG (8). More recent studies showed promise for mRNA vaccines against group A and B *Streptococcus*, *Bordetella pertussis*, *Yersinia pestis*, *Chlamydia trachomatis*, and *Borrelia burgdorferi* (9–14). The mRNA platform has since been extended to a broad range of extracellular, toxin-driven, and multidrug-resistant bacteria, including *Clostridioides difficile*, *Pseudomonas aeruginosa*, *Klebsiella pneumoniae*, *Acinetobacter baumannii*, and *Streptococcus pneumoniae*, and has frequently outperformed matched protein subunit-based vaccine comparators (16–18, 19). The first bacterial mRNA candidates have now entered clinical evaluation, with mRNA–LNP vaccines against *Borrelia burgdorferi* (OspA) and *M. tuberculosis* advancing to early-phase human trials (NCT05975099 and NCT05537038, respectively) (5).

To address the urgent need for a gonococcal vaccine, we, for the first time, evaluated an LNP-encapsulated mRNA encoding the highly conserved antigen BamA as a preclinical candidate. According to the World Health Organization (WHO), gonorrhea affected 82.4 million individuals worldwide (19, 20), and *Neisseria gonorrhoeae* (*Ng*) is classified as a high-priority pathogen for the development of new interventions in the WHO Bacterial Priority Pathogens List (21). Left untreated, gonorrhea can cause severe sequelae, including infertility, pelvic inflammatory disease, preterm birth, and neonatal blindness (22). Surveillance efforts have documented a rise in multidrug-resistant *Ng* isolates over recent decades. Owing to rising antibiotic resistance, azithromycin was removed from the recommended dual-antibiotic regimen in 2020, leaving ceftriaxone as the only treatment for uncomplicated gonorrhea (23). Recently, two oral therapies, zoliflodacin and gepotidacin, were approved by FDA to treat uncomplicated urogenital gonorrhea (24). Diverse vaccine platforms have been explored against *Ng*, including partially inactivated whole cells pooled from two to three strains, pilus protein, outer membrane and outer membrane vesicles (OMVs), recombinant proteins formulated with different adjuvants, the lipooligosaccharide epitope 2C7, DNA encoding PorB, virus replicon particles, bacterial ghosts, and Tag/Catcher capsid virus-like particles (22, 25–27). A potential breakthrough came from an observational case-control study reporting 31% efficacy of the *N. meningitidis* serogroup B OMV vaccine MeNZB against *Ng* infection (28). Retrospective studies of the derivative meningococcal vaccine 4CMenB (Bexsero; GSK), which contains MeNZB OMVs plus four purified recombinant antigens, have further confirmed reduced rates of gonococcal infection, with an estimated effectiveness of 30–46% (16, 22, 29). In response, the United Kingdom implemented a targeted 4CMenB program for high-risk individuals in August 2025 (30).

Immunization of mice with 4CMenB vaccine resulted in accelerated clearance of *Ng* and the murine antibodies cross-reacted with several gonococcal antigens, including BamA, suggesting BamA as one of the cross-protecting antigens (31). Previous to these findings, we have identified BamA as a desirable target for gonococcal vaccine development by proteomics and bioinformatics-based vaccinology approaches (32–35). BamA is a β-barrel outer membrane protein (OMP) and the central component of the β- barrel assembly machine, BAM (36). This heteromeric complex, in addition to BamA, contains a variable number of outer membrane lipoproteins (BamB-F) that work together to drive the assembly and insertion of the β-barrel OMPs in all Gram-negative bacteria (36, 37). BamA is ubiquitously expressed by geographically, temporally, and genetically diverse gonococcal isolates and is an essential protein for *Ng* viability (33, 34). The structure of full-length *Ng* includes the protein extracellular loops that form a dome over a large C-terminal β-barrel membrane domain and five periplasmic polypeptide-transport-associated (POTRA) domains located near the barrel pore. Bioinformatic analyses of the occurrence of single nucleotide- and single amino acid polymorphisms (SAAPs) demonstrated that BamA was one of the most highly conserved antigens among 34 examined immunogens in over 5,000 *Ng* clinical isolates globally. Only two low-prevalence single amino acid polymorphisms (1-10% isolates) were identified in BamA surface-exposed loops, while the most SAAPs were mapped to the β-barrel and POTRA domains that are buried in the outer membrane and periplasm, respectively (32). Despite these desirable characteristics, purifying BamA in its native conformation in amounts sufficient for preclinical studies and later-stage vaccine production remains challenging (33, 36). Herein, we explored an mRNA platform that circumvents these difficulties and formulated BamA mRNA in lipid nanoparticles (LNPs), which were administered to female BALB/c mice via intramuscular or intranasal immunization and were followed by intravaginal challenge with two diverse *Ng* isolates, FA1090 and WHO X.

## Materials and Methods

### BamA Antigenicity and Epitope Prediction

The amino acid sequence of the BamA protein of the *Ng* FA1090 was downloaded from the UniProt database (UniProt ID: Q5F5W8), and the 3D structure of the antigen (PDB ID: 4K3B) was obtained from the Protein Data Bank (36). To evaluate the immunogenic potential and antigenicity of the Q5F5W8, we employed VaxiJen, a web-based server for predicting protective antigens for subunit vaccines (38). VaxiJen predicts the likelihood that a protein is an immunogen based on its physicochemical properties, using three distinct models to assess immunogenic potential, and provides a probability score for its immunogenicity and antigenicity.

Linear and conformational (discontinuous) B-cell epitopes were predicted using the ElliPro server (http://tools.iedb.org/ellipro/), which implements Thornton’s method for identifying protruding surface regions using the Protrusion Index (PI) and a residue clustering algorithm to predict antibody-accessible epitopes (39). Default parameters were used, with a minimum score threshold of 0.8 and a maximum distance of 6 Å between residues for discontinuous epitope clustering. All predicted linear and discontinuous epitopes were ranked by their PI scores, and the top-scoring epitopes were selected for structural mapping. Residue numbering was based on the full-length precursor (initiator methionine at position 1) to maintain consistency with ElliPro. We predicted high-affinity MHC class I (CD8+) and class II (CD4+) T-cell epitopes within BamA. For MHC class I, 9-mer peptides spanning the entire BamA sequence were evaluated for binding to a panel of 10 common human HLA-A, HLA-B, and HLA-C alleles using NetMHCpan-4.1 (http://services.healthtech.dtu.dk/service.php?NetMHCpan-4.1), which integrates both peptide-MHC binding affinity and eluted ligand data. For MHC class II, NetMHCIIpan-4.0 (http://services.healthtech.dtu.dk/service.php?NetMHCIIpan-4.0) was employed to predict binding (40). Peptides with a predicted % rank of <1.0 (strong binders) or <2.0 (weak binders) were considered high-affinity binders. Peptides with scores >0.9 and promiscuous binding across multiple HLA supertypes were prioritized for downstream analysis.

### Bacterial Strains and Growth Conditions

*Ng* strains, FA1090 and the 2016 WHO reference strains (41, 42), were grown on gonococcal base agar (Becton, Dickinson and Company) supplemented with Kellogg’s supplement I and 12.5 μM ferric nitrate at 37 °C with 7% carbon dioxide for up to 20 h and in gonococcal base liquid medium (Becton, Dickinson and Company) supplemented with Kellogg’s supplement I and 12.5 μM ferric nitrate (27, 43).

*Escherichia coli* BL21 (DE3) bearing a plasmid overproducing *Ng* FA1090 BamA protein(34) was maintained on Luria-Bertani agar and cultured in Luria-Bertani broth (Becton, Dickinson and Company), supplemented with carbenicillin at 50 μg/mL. Overproduction of BamA was induced with 1 mM isopropyl-β-D-galactopyranoside (IPTG) after the bacteria reached an OD_600_ of 0.5 and were grown for an additional 3 h at 37 °C with aeration (34).

### DNA Manipulations

Cloning procedures and oligonucleotides were designed using SnapGene software version 2.8 (GSL Biotech LLC). Q5 High-Fidelity DNA polymerase, DNA ligase, and NEBuilder HiFi DNA Assembly Master Mix were obtained from New England Biolabs (NEB). The *Ng* FA1090 *bamA* gene lacking the amino acids encoding BamA signal peptide was amplified using purified genomic DNA from Ng FA1090 and primers BamAnoSP-f, AAGCTGGCTAGCCACCATGGACTTCACCATCCAAGACATC and BamAnoSP-r, GCTGATCAGCGGGTTTAAACTTAGAACGTCGTGCCGAG. The *bamA* gene with the murine IgGκ signal peptide (mIgGκSP) was amplified using primers BamAwSP-f, TTCCAGGTTCCACTGGTGACGACTTCACCATCCAAGAC and BamAwSP-r, GCTGATCAGCGGGTTTAAACTTAGAACGTCGTGCCGAG. The resulting PCR products were digested with EagI (NEB), ligated into similarly digested pSecTag2a (Invitrogen), and introduced into competent *E. coli* BL21(DE3). DNA constructs were verified by Sanger Sequencing at the Center for Quantitative Life Sciences at Oregon State University.

### RNA Design and Synthesis

The BamA mRNA construct was modified with 5moU to reduce the transcript’s immunogenicity. In one of the two tested mRNA constructs, the murine IgGκ signal peptide preceded the *bamA* coding sequence. The BamA mRNA was treated with a phosphatase to remove immunogenic 5′-triphosphates and formulated in 1 mM sodium citrate buffer at pH 6.4. BamA mRNA and Enhanced Green Fluorescent Protein (EGFP) mRNA were purchased from Trilink Biotechnologies.

### Lipid Nanoparticle Formulation and Characterization

The lipid nanoparticle (LNP) formulation consisted of SM-102, cholesterol, 1,2- distearoyl-sn-glycero-3-phosphocholine (DSPC), and dimyristoyl-rac-glycero-3- methoxypolyethylene glycol-2000 (DMG-PEG_2k_) at a molar ratio of 50:38.5:10:1.5 (the standard lipid composition), with the mRNA present at an N/P ratio of 5.30. Cholesterol and DMG-PEG_2k_ were purchased from Sigma-Aldrich and NOF America, respectively. SM-102 was purchased from Cayman Chemicals. DSPC was obtained from Avanti Polar Lipids, Inc. The LNPs were prepared by combining the aqueous phase containing mRNA in 50 mM citrate buffer, pH 4.0, with the ethanolic phase containing lipids at a 3:1 ratio using a microfluidic mixer (Ignite, Precision Nano Systems). The two phases were injected into the microfluidic device at a combined volumetric flow rate of 9 mL/min. Formulations were then diluted with Phosphate-Buffered Saline (PBS, pH 7.2) and dialyzed for 4 h at room temperature against sterile PBS, followed by overnight dialysis at 4°C in 10 kDa Slide-a-Lyzer G2 cassettes (Thermo Fisher). They were then concentrated using Amicon Ultra Centrifugal Filters (EMD Millipore), characterized, and stored at 4°C until use.

LNPs were characterized for particle size and polydispersity index (PDI) using dynamic light scattering using Stunner (Unchained labs). mRNA encapsulation efficiency and concentration were evaluated by the modified Quant-iT Ribogreen RNA reagent assay (Life Technologies). LNP sample or PBS (negative control) was diluted into Tris EDTA (TE) buffer to prepare the working solution. Aliquots of each LNP working solution were further diluted with either 1:1 TE buffer to measure unencapsulated free mRNA or 1:1 TE buffer supplemented with 2% Triton-X to measure total mRNA, which includes both free unencapsulated mRNA and LNP-encapsulated mRNA. Samples were prepared in triplicate. Quanti-iT™ RiboGreen RNA reagent was added to each sample, and the fluorescent signal was quantified (Tecan i-Control v. 3.8.2.0, Tecan Infinite M200 Pro Multimode Plate Reader). The LNPs were diluted to the desired concentration with sterile, endotoxin-free PBS for administration in mice.

### Cell Line Transfection and Fractionation

Human Embryonic Kidney (HEK) 293 cells were cultured in Dulbecco’s Modified Eagle’s Medium (DMEM) supplemented with 10% Fetal Bovine Serum (FBS) in an incubator at 37°C with 5% carbon dioxide. HEK293 cells were treated with lipofectamine alone as a mock transfection, an empty pSecTag2A vector, a pSecTag2A vector (Thermo Scientific) with the *bamA* gene missing signal peptide, as well as a pSecTag2A vector with the signal peptide *mIgGκSP* fused to *bamA*. The pSectag2A vectors contain a Zeocin resistance gene for selection in mammalian cells. Cells were transfected with Lipofectamine and treated with Zeocin as a selective antibiotic after transfection. Cells and their respective culture supernatants from each group treatment were prepared to evaluate *bamA* expression and BamA localization as follows. HEK293 cells were harvested by centrifugation at 1,000×*g* at 10°C for 15 min (44) and lysed in buffer consisting of 50 mM Tris HCl, 150 mM NaCl, 1.0% (v/v) NP-40, 0.5% (w/v) sodium deoxycholate, 1 mM EDTA, 0.1% (w/v) SDS, pH 7.4, at 4°C before boiling for 4 min at 100°C. Cells were centrifuged, sonicated for four cycles at 10 sec/cycle, and centrifuged at 1,041×*g* for 3 min at 25°C before the soluble fraction of cell lysates was subjected to protein quantification by the DC Protein Assay (Bio-Rad). To concentrate the proteins in the culture supernatants, samples were precipitated using the pyrogallol red-molybdate- methanol method (45). Proteins were quantified using the DC Protein Assay (Bio-Rad).

### SDS-PAGE and Immunoblotting

To assess the presence of *Ng* BamA in HEK293 cells and in their culture supernatants, samples prepared as outlined above were normalized by protein amounts and separated by sodium dodecyl sulfate polyacrylamide gel electrophoresis (SDS-PAGE). Whole-cell lysates of the *E. coli* pBamA and the 2016 *Ng* WHO reference strains were normalized to optical density (OD_600_) and lysed with Laemmli Sample Buffer (Bio-Rad) (27, 33, 46, 47). For all samples, proteins were separated in Criterion TGX gels (Bio-Rad) before transfer to nitrocellulose membranes using the TransBlot Turbo (Bio-Rad).

To detect BamA in HEK293 cells and their culture supernatants, membranes were blocked in PBS with 0.1% Tween 20 (PBST) in 5% non-fat milk, washed in PBST, and probed with polyclonal BamA rabbit antisera (1:20,000) for 1 h in PBST with 5% milk. Membranes were then washed with PBST and incubated for 1 h with 1:10,000 diluted horseradish peroxidase–conjugated goat anti-rabbit secondary antibodies (Southern Biotech) and developed as described below.

Detection of BamA-specific serum and vaginal IgG and IgA was performed as described (27, 48). Briefly, membranes were blocked at 4°C overnight in PBST and 5% milk. After washing with PBST, the membranes were incubated for 1 h at room temperature with pooled mouse sera diluted 1:5,000. Pooled vaginal lavages were diluted 1:50 in PBST, and 5% milk and membranes were incubated in the solution at 4°C overnight. Subsequently, the blots were probed with goat anti-mouse IgG (BioRad) or IgA (Southern Biotech) conjugated to HRP (1:10,000 dilution) as described previously (27, 46, 48). Cross-reacting proteins were detected using ECL Prime (Amersham) and ImageQuant^TM^ LAS 4000 (GE Healthcare) with the ChemiDoc MP Imaging System (Bio-Rad).

### Preparation of naturally secreted nOMVs

Naturally secreted nOMVs from the *Ng* FA1090 strain were harvested during the stationary phase of growth, prepared as previously described (34, 49), and reconstituted in PBS at 4°C overnight. Protein concentrations were quantified using the DC Protein Assay (Bio-Rad).

### Immunization and Challenge Experiments

To minimize bias, researchers were blinded to all immunization and challenge experiments. In the initial intramuscular (IM) immunization experiment (pilot IM study), four-week-old female BALB/c mice (Charles River) were divided into three cohorts (*n*= 6- 8 mice/cohort) and immunized in three doses (10 µg/dose) and three weeks apart with endotoxin-free PBS, EGFP mRNA (Trilink BioTechnologies), or BamA mRNA (Trilink BioTechnologies), both encapsulated in LNPs. Retroorbital blood and vaginal lavage collections were performed on days 31, 52, and 63 (relative to the prime immunization, designated as day 0). To collect vaginal secretions, 33 µL of endotoxin-free PBS was pipetted three times in the mouse vaginal canal. For retro-orbital bleeding, 100-150 µL of blood was collected from the retro-orbital sinus via capillary tubes (Kimble) before mice were treated with 0.5% proparacaine (27, 46, 48). Blood and vaginal lavage collections were centrifuged at 4°C for 10 and 5 min, respectively, before the isolated serum and vaginal lavage samples were stored at -80°C.

Two independent immunization/challenge experiments with BamA mRNA-LNPs- based vaccines were performed using four-week-old female BALB/c mice (Charles River) (*n*= 15-25 mice/cohort). For the first experiment, mice were IM-immunized with either PBS, 10 µg of EGFP mRNA, or 10 µg of BamA mRNA, all in LNP, on days 0, 21, and 42. Retro-orbital bleeds were collected on days 31 and 52, whereas vaginal lavages were collected on day 31. Three weeks after the third vaccine dose, mice in diestrus stage of the estrous cycle were treated with a subcutaneous implant of slow-release 6.5mg 17-β estradiol pellet (Innovative Research of America) and a combination of 2.4 mg streptomycin sulfate (VWR) and 0.4 mg vancomycin hydrochloride (VWR) intraperitoneally and 0.04 g trimethoprim sulfate (MP Biomedicals) per 100 mL volume in given drinking water (50, 51). Two days after pellet implantation, mice were inoculated intravaginally with 10^6^ CFU of piliated *Ng* FA1090 as described (46, 50–52). Vaginal swabs for CFU enumeration were collected using sterile rayon-tipped applicators (Puritan, 25-800 1PD 50) on days 1, 3, 5, and 7 post-infection. Dilutions of vaginal swab specimens in GC broth containing 1 % (w/v) saponin were spread using the Eddy Jet 2W Spiral Plater (IUL Instruments) and maintained on GC agar supplemented with VCNT Inhibitor (Becton, Dickinson and Company) of selective antibiotics 1.5 mg vancomycin, 3.75 mg colistin, 6,250 U nystatin, 2.50 mg trimethoprim sulfate, and 100 mg streptomycin sulfate (VWR) for 30 h in 37°C with 7% carbon dioxide. The lower limit of detection was 200 colony-forming units (CFUs)/mL per plate, as determined using the Automated Colony Counter SphereFlash (Neutec Group Inc).

In the second immunization/challenge study, mice were immunized intranasally (IN) on days 0, 21, and 42 using PBS, 15 µg CpG oligodeoxynucleotides (ODN 2395; Invivogen), 10 µg BamA mRNA LNPs, and 10 µg BamA mRNA combined with 15 µg CpG 2395 ODN and encapsulated in LNPs (BamA mRNA LNPs+CpG). Blood and vaginal lavage samples were collected as described above. Mice were administered estradiol through subcutaneous injections of 0.5 mg Premarin (Pfizer) two days before infection, on the day of infection, and two days after infection (53–55). Mice were inoculated intravaginally with piliated Ng WHO X at 10^6^ CFUs. Vaginal swabs were collected on days 1, 3, 5, 7, 9, 11, and 13 post-infection and plated for CFUs enumeration as described above. Investigators were blinded during all immunization and challenge experiments.

### Ethics Statement

Animal work was conducted in accordance with the standards of the Association for Assessment and Accreditation of Laboratory Animal Care and performed under the Oregon State University Institutional Animal Care and Use Committee. Animal facilities adhere to the NIH Guide for the Care and Use of Laboratory Animals (Publication No. 85- 23) and function under the supervision of a full-time attending veterinarian. Mice were monitored throughout the study and euthanized at experimental endpoints by isoflurane inhalation using a regulated compressed-gas system, followed by cervical dislocation to confirm death, in accordance with institutional guidelines and the AVMA Guidelines for the Euthanasia of Animals (56).

### Enzyme-Linked Immunosorbent Assay

Antibody types and endpoint titers in mouse sera and vaginal lavages were analyzed through an Enzyme-linked Immunosorbent Assay (ELISA) as previously described (27, 46, 48) with the following modifications. The antigen coating consisted of *Ng* FA1090 nOMVs, prepared as described above, at 50 ng/well in coating buffer containing 1 L of dd water, 1.5 g of anhydrous sodium carbonate, and 2.93 g of anhydrous sodium bicarbonate at pH 9.6. 96-well clear polystyrene high-binding round U-bottom plates (Greiner Bio-One) were coated with antigen coating buffer and stored overnight at 4°C, then blocked for 1 h at room temperature with 1:4-diluted Block Ace (Bio-Rad). Mouse sera were diluted with PBST at 1:243 to obtain 900 μL volumes, which were dispensed into the first column of Nunc 96-well polypropylene DeepWell Sample Plates (Thermo Scientific). All other columns of the DeepWell Sample Plates received 600 μL of PBST. Three-fold serial dilutions of the 1:243 mouse serum solutions were performed horizontally across the plate from the first to the tenth column using the Integra Viaflo 96 (Agilent Technologies). The eleventh and twelfth columns of the DeepWell Sample Plates contained only PBST, using the eleventh column to determine the background absorbance. The serially diluted samples were dispensed from the DeepWell Samples

Plates into the 96-well U-bottom plates (Greiner Bio-One) at 100 μL per well using the Integra Viaflo 96. Individual mouse vaginal lavage samples were diluted with PBST at 1:81 in 450 μL, which were dispensed into the first column of the DeepWell Sample Plates. All other columns in the DeepWell received 300 μL PBST. The 1:81 diluted vaginal lavage samples in the first column underwent three-fold serial dilution horizontally across the plate from the first column to the tenth column. The eleventh and twelfth columns contained only PBST. The Integra ViaFlo 96 was used to dispense 100 μL per well from the DeepWell Sample Plates into the 96-well U-bottom plates (Greiner Bio-One). Following overnight incubation with serially diluted sera or vaginal lavages at 4°C, plates were washed using the Biotek 405 LS microplate washer (Agilent Technologies) three times with PBST before 1 h room temperature incubation with 1:5,000 Horseradish peroxidase–conjugated goat anti-mouse secondary antibodies Ig, IgA, IgG1, IgG2A, and IgG3 (Southern Biotech) as described (27). Plates were washed three times with PBST before adding 1-Step Ultra TMB Substrate (Thermo Scientific) to all wells, followed by adding 0.2 M sulfuric acid to stop the reaction. Absorbance was recorded at an optical density of 450 nm using the Biotek Synergy HT Plate Reader (Agilent Technologies).

### Serum Bactericidal Assay

The human complement-dependent serum bactericidal assay (SBA) was performed as previously described (27, 48). Murine sera from immunized and control groups were pooled, heat-inactivated at 56°C for 30 min and serially diluted two-fold in Hanks’ Balanced Salt Solution (HBSS) across a range of 1:64 to 1:32,768. Non-piliated *Ng* WHO X (1×10³ CFU in 40 µL HBSS) were added to wells containing diluted test sera and incubated for 15 min at 37°C. Subsequently, 10 µL of either IgG/IgM-depleted normal human serum (NHS; Pel-Freeze Biologicals; Cat. # 34010-10, Lot 34943) or heat-inactivated NHS (HI-NHS; Cat. #34020-10, Lot 13844) was added as a 10% (v/v) complement source, and reactions were incubated for 1 h at 37°C. A single lot of each serum was used across all assays to minimize complement-related variability. Bactericidal activity was assessed by spot inoculating 5 µL aliquots and their corresponding 10-fold dilutions onto chocolate agar, followed by overnight incubation at 37°C for CFU enumeration. Assay controls included: *Ng* + NHS, *Ng* + HI-NHS, *Ng* + test sera, and *Ng* alone. Percent killing at each dilution was calculated by comparing CFUs recovered from samples containing test sera with active NHS to CFUs recovered from matched vials containing test sera with HI-NHS. SBA titers were defined as the reciprocal of the highest serum dilution yielding ≥50% killing. All assays were performed in three biological replicates.

### Statistical analysis

Statistical analyses were performed using GraphPad Prism (version 11; GraphPad Software). *Ng* clearance rates were compared between groups using the Kaplan-Meier method with the log-rank (Mantel-Cox) test. Bacterial burden at each time point was compared using multiple Mann-Whitney tests with FDR correction (Benjamini-Hochberg method). The area under the curve (AUC) of log_10_-transformed CFU/mL over the seven-day infection period was calculated using the trapezoid rule and compared between groups by Mann-Whitney test. ELISA antibody titers were log10-transformed prior to analysis (27, 46, 48). Data were assessed for normality using the Shapiro-Wilk test. Group differences were evaluated using the Kruskal-Wallis test, followed by Dunn’s post hoc test with a Bonferroni correction for multiple comparisons. Statistical significance is denoted as follows: \**p* < 0.05, \*\**p* < 0.01, \*\*\**p* < 0.001, \*\*\*\**p* < 0.0001. The IgG2a:IgG1 ratio was calculated for each mouse as the difference between log_10_-transformed IgG2a and IgG1 AUC values, and the differences were assessed by a Kruskal-Wallis test with Dunn’s multiple comparisons post hoc test and Bonferroni correction. Sample sizes, replicates, and specific tests used are indicated in the corresponding figure legends.

### Visualization and illustrations

Data graphs were generated using GraphPad Prism 11. Schematic illustrations were created with BioRender.com and Adobe Illustrator.

## DATA AVAILABILITY

All data supporting the findings of this study are available within the article and its Supplementary Information files. The raw data can be obtained from the corresponding author upon request.

## Results

### BamA contains B-cell epitopes in POTRA and β-barrel domains and broadly distributed T-cell epitopes

To further assess the potential of *Ng* FA1090 BamA as a gonococcal vaccine candidate, we used the VaxiJen, ElliPro, NetMHCpan-4.1, and NetMHCpan-4.0 bioinformatic tools. Three VaxiJen models independently predicted, with 100% probability, that BamA was immunogenic, with an antigenicity score of 0.6518. ElliPro identified four linear and three discontinuous B-cell epitopes with a stringent score threshold of 0.8 (**Table 1**). The two highest-scoring linear epitopes mapped to POTRA domain 1; residues 23–61, score 0.934, and residues 73–89, score 0.895 (**Fig. 1**). Additional linear epitopes mapped to POTRA 3 (residues 261–282, score 0.859) and the POTRA 4-5 junction (residues 327–342, score 0.870). The top discontinuous epitope (score 0.918) clustered across POTRA 1 and POTRA 2, while the remaining discontinuous epitopes localized to POTRA 2–4 (score 0.853) and the transmembrane β- barrel strands (score 0.840). None of the predicted B-cell epitopes localized to the surface-exposed loops L1, L5, or L8 (**Fig. 1)**.

**Figure 1.**
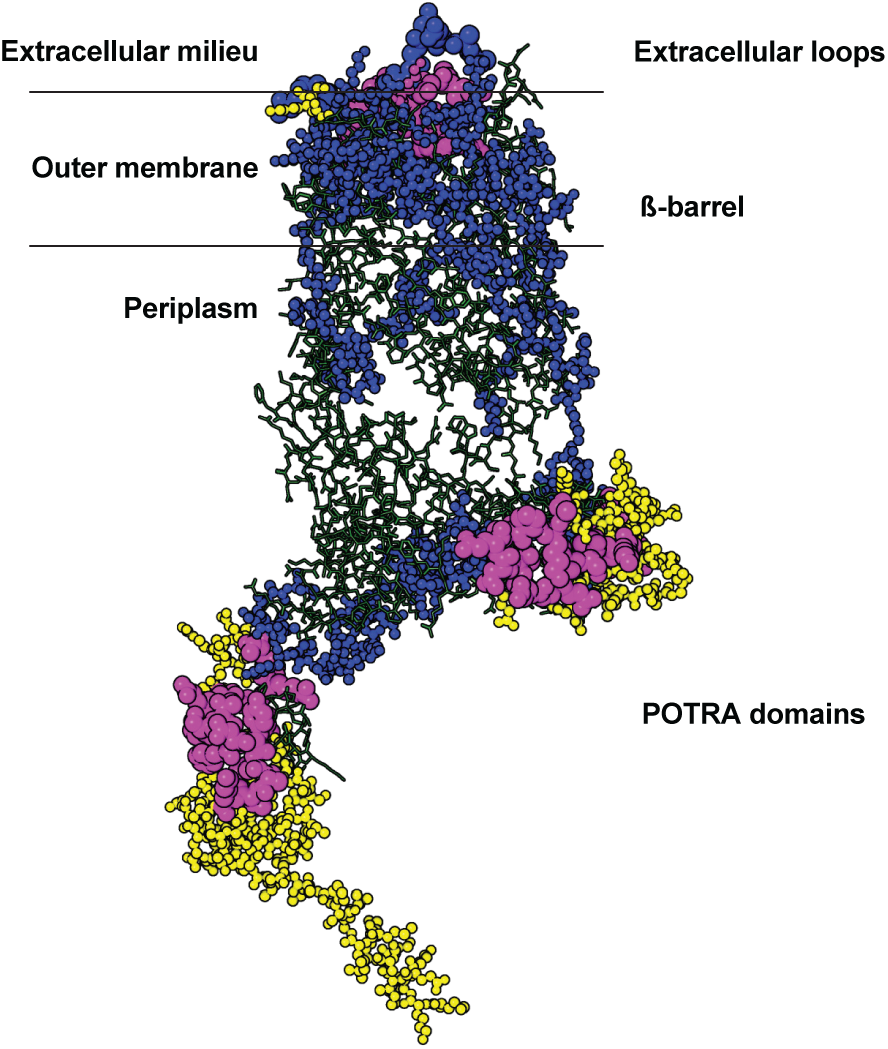
Predicted B-cell epitopes of *Neisseria gonorrhoeae* BamA cluster in the periplasmic POTRA and membrane-embedded β-barrel domains, whereas high-affinity MHC class I T-cell epitopes localize to the surface-exposed extracellular loops. Surface representation of the full-length *N. gonorrhoeae* FA1090 BamA structure (PDB 4K3B), shown in green. Predicted B-cell epitopes are represented as yellow spheres and predicted T-cell epitopes as blue spheres, indicating their distribution across the protein structure. Residues at which B-cell and T-cell epitopes overlap are highlighted as magenta spheres. B-cell epitopes were predicted with ElliPro using a minimum Protrusion Index score of 0.8 and a maximum inter-residue distance of 6 Å; MHC class I T-cell epitopes were predicted with NetMHCpan-4.1 across a panel of 10 common human HLA-A, HLA-B, and HLA-C alleles.

**Table 1.**
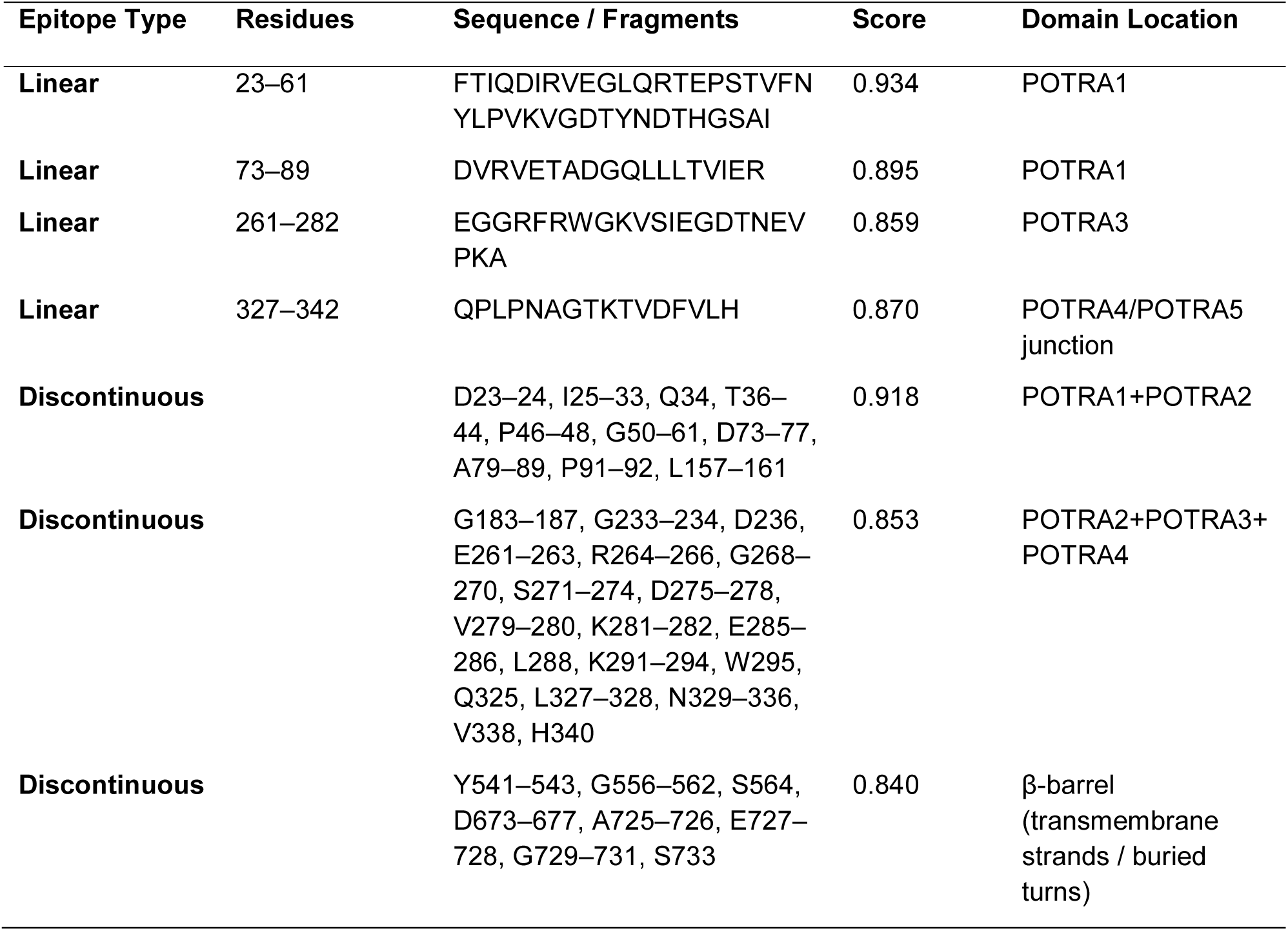
Predicted Linear B-Cell Epitopes in the BamA Protein of *Neisseria gonorrhoeae*.

To examine the global human population coverage, we have also predicted MHC class I and class II T-cell epitopes using a panel of 10 common HLA class I and class II alleles (**Table 2**). We identified 28 and 8 high-affinity MHC-I and MHC-II epitopes, respectively, with scores exceeding 0.9 (**Table 3**). Importantly, MHC-I epitopes mapped to the BamA extracellular loops L1, L5, and L8 with scores of 0.953, 0.817, and 0.981, respectively (**Fig. 1**). The top-scoring MHC-I epitope, YSATHNQTW (score 0.991), bound HLA-B*57:01 and localized to a transmembrane β-strand. Several high-affinity epitopes mapped to the periplasmic POTRA domains (e.g., LTEGGIWTW and KLNIQITPK, both in POTRA 2, with scores of 0.976 and 0.955, respectively, and AELEKLLTM in POTRA 4, score 0.887, confirming efficient intracellular processing of these BamA regions.

**Table 2:** HLA Alleles Used for T-Cell Epitope Prediction.

| HLA | Allele | Supertype | Global Prevalence / Population Relevance |
| --- | --- | --- | --- |
| HLA Class I | HLA-A*01:01 | A1 | Present in European populations |
|  | HLA-A*02:01 | A2 | Very high (Europe, Americas, Asia) |
|  | HLA-A*03:01 | A3 | Common in Caucasians |
|  | HLA-A*11:01 | A3 | High in East and Southeast Asians |
|  | HLA-A*24:02 | A24 | High in Asian and Oceanian populations |
|  | HLA-B*07:02 | B7 | High in Europeans |
|  | HLA-B*08:01 | B8 | Common in European populations |
|  | HLA-B*15:01 | B62 | Frequent in Asians and Latinos |
|  | HLA-B*44:02 | B44 | Common in Europeans |
|  | HLA-B*57:01 | B58 | Notable for strong immune responses (e.g., HIV control) |
| HLA Class II | HLA-DRB1*01:01 | DR | High in Europeans |
|  | HLA-DRB1*03:01 | DR | Common globally |
|  | HLA-DRB1*04:01 | DR | Prevalent in Europe and North America |
|  | HLA-DRB1*07:01 | DR | Broadly distributed worldwide |
|  | HLA-DRB1*09:01 | DR | Frequent in East Asian populations |
|  | HLA-DRB1*11:01 | DR | Common in diverse global populations |
|  | HLA-DRB1*13:01 | DR | Present across Africa, Asia, and Europe |
|  | HLA-DRB1*15:01 | DR | Very common globally |
|  | HLA-DQA1*01:02 /<br>DQB1*06:02 | DQ | Found in multiple ethnic groups |
|  | HLA-DPA1*01:03 /<br>DPB1*04:01 | DP | Widely represented across populations |

**Table 3:**
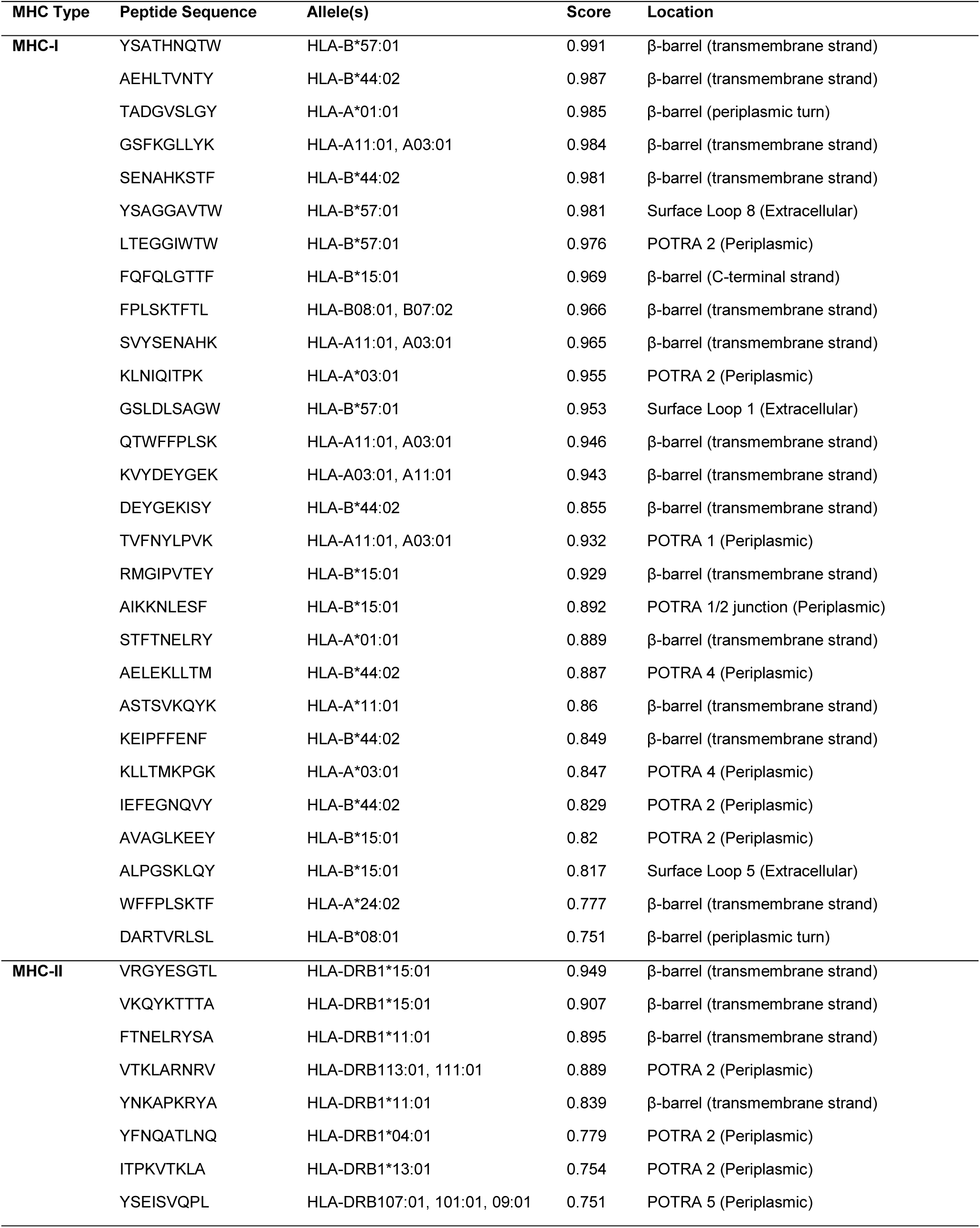
Predicted T-cell epitopes within BamA with Domain Location.

### Recombinant *Ng* BamA is expressed in mammalian cells and released into the extracellular milieu

To evaluate if *Ng* BamA is translated in mammalian cells, we created two constructs in the pSecTag2A mammalian expression vector: one with the *bamA* gene lacking a cognate signal peptide and another with a truncated *bamA* preceded by the signal sequence of the murine immunoglobulin (Ig)κ (mIgGκSP; **Fig. S1A**).

We then examined the presence of the *Ng* BamA protein in HEK293 cells and in their tissue culture medium using polyclonal rabbit antisera raised against recombinant *Ng* BamA (34). The immunoblotting showed that the BamA antisera readily cross-reacted with a major protein band migrating at ∼88 kDa and ∼100 kDa, corresponding to the expected molecular masses of BamA lacking the signal peptide and containing mIgGκSP, respectively, in both cells and supernatants of HEK293 cells carrying pSecTag2A-*bamA* or pSecTag2A-Igκ*bamA*. We observed a slight increase in the amount of BamA containing the murine Igκ light chain leader sequence, suggesting that the presence of a murine signal peptide may increase the expression or stability of BamA in eukaryotic cells (**Figs. S1B** and **S1C**, respectively).

### BamA mRNA design and LNP characterization

The presence of murine Igκ enhanced antigen expression and processing in host cells, thereby augmenting vaccine efficacy (57–60). Thus, following successful expression of BamA in HEK293 cells, a non-amplifying mRNA (NRM) construct was designed using the BamA coding sequence with an N-terminal mIgGκSP tag and encapsulated in LNPs (**Fig. 2A**). As controls for the immunization/challenge experiments, we also included EGFP and a CpG adjuvant. The formulated LNPs ranged in size from 72.40 ± 0.68 nm for EGFP mRNA LNPs to 68.84 ± 7.94 nm for CpG LNPs, 71.81 ± 6.83 nm for BamA mRNA LNPs, and 82.15 ± 6.31 nm for BamA+CpG LNPs (**Fig. 2B**). The particles showed mRNA encapsulation higher than 95%, reported as 97.87 ± 0.32% for EGFP mRNA LNPs, 98.73 ± 0.12% for CpG LNPs, 98.62 ± 0.51% for BamA mRNA, and 98.80 ± 0.60% for BamA+CpG LNPs (**Fig. 2C**). When the LNPs were assessed for charge density at the LNP surface, the zeta potential values were determined to be -4.00 ± 0.75 mV for EGFP mRNA LNPs, -2.85 ± 2.13 mV for CpG LNPs, -3.31 ± 2.11 mV for BamA mRNA LNPs, and -3.38 ± 2.15 mV for BamA+CpG LNPs (**Fig. 2D**). The polydispersity index (PDI) indicating the heterogeneity for the size of LNPs were reported over the following ranges: 0.09 ± 0.01 for EGFP mRNA LNPs, 0.07 ± 0.05 for CpG LNPs, 0.13 ± 0.05 for BamA mRNA LNPs, and 0.27 ± 0.03 for BamA+CpG LNPs (**Fig. 2E**). The higher PDI value for co-encapsulated BamA+CpG LNPs maybe reflect two combined factors; the centrifugation-mediated concentration required to reach the target dose within the small intranasal volumes tolerated by mice, and the altered physicochemical environment introduced by CpG co-encapsulation (61).

**Figure 2.**
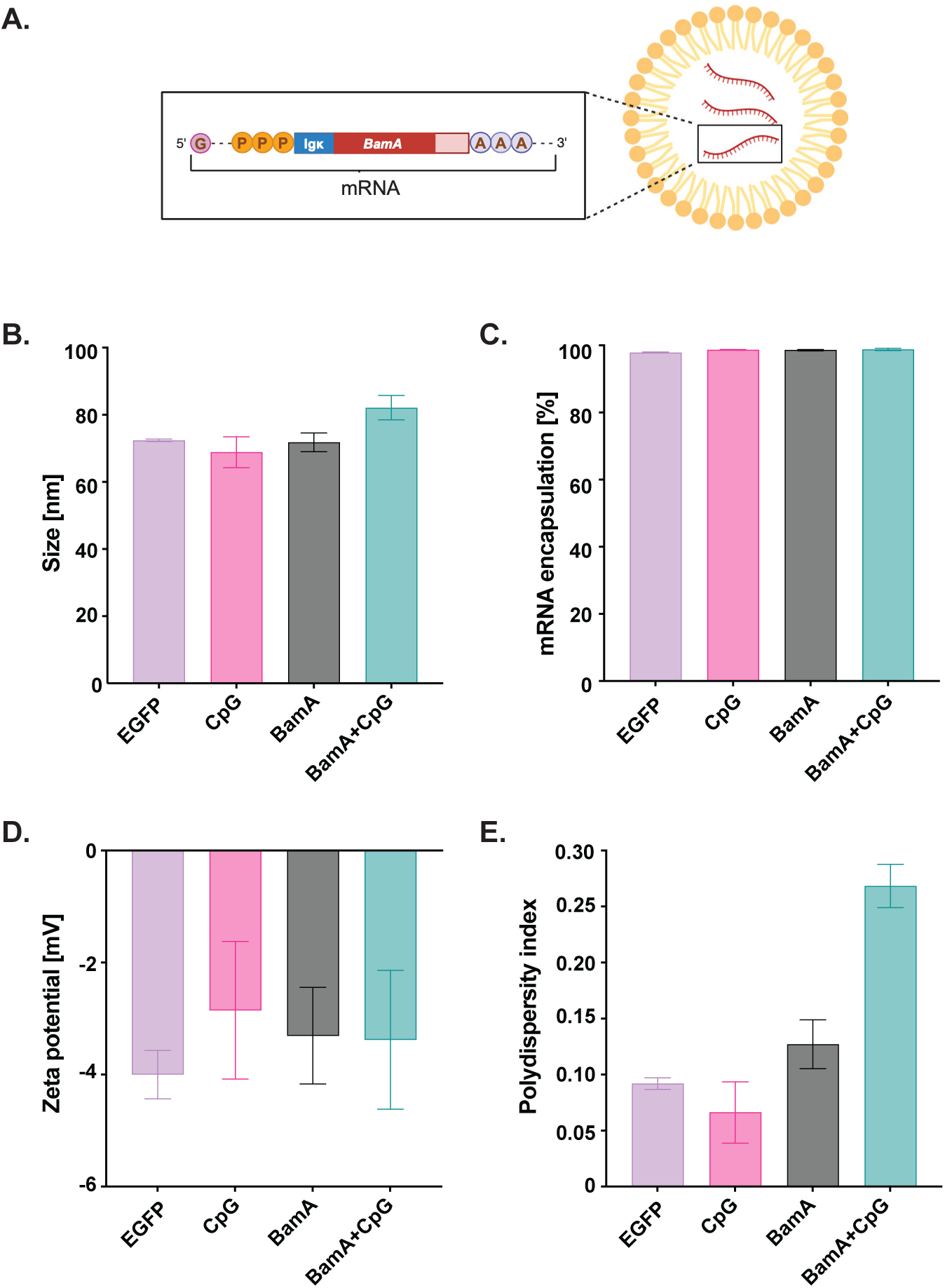
Design and physicochemical characterization of BamA mRNA–lipid nanoparticles (LNPs). **A**, Schematic of the non-amplifying BamA mRNA construct carrying the mouse IgGκ signal peptide upstream of the bamA coding sequence and 5- methoxyuridine (5moU) nucleoside modification, encapsulated in SM-102-based LNPs. Hydrodynamic diameter (**B**), mRNA encapsulation efficiency (**C**), zeta potential (**D**), and polydispersity index (**E**) of LNPs formulated with EGFP mRNA, CpG, BamA mRNA, or BamA + CpG mRNA. Bars show mean ± SD from three independent formulations.

### Experimental outline of preclinical BamA mRNA-LNP studies

To assess the immune responses elicited by BamA mRNA LNPs, we conducted three independent studies: pilot IM immunization (**Fig. 3A**), IM immunization followed by challenge with Ng FA1090 (**Fig. 3B**), and IN immunization and gonococcal challenge with Ng WHO X (**Fig. 3C**). The IM route drives rapid antigen uptake, accommodates a wide range of injection volumes, and supports efficient antigen presentation near the injection site (62). IM delivery is also the most widely used route of vaccination in clinical practice, offering patient familiarity and translatability to human clinical trials, as demonstrated by the COVID-19 mRNA vaccines (63). It sustains protein expression at the injection site for up to 10 days after LNP delivery, longer than other routes (64). We have also turned to a mucosal route to induce antigen-specific antibodies at the site of *Ng* infection. An IN immunization with mRNA encoding the Hsp65 protein was protective against a TB challenge in mice (7). The IN administration of nOMVs and protein subunit vaccines resulted in accelerated *Ng* clearance (22, 25, 46, 65–68), and prior work also suggested that a Th1 response may be required to elicit protective immune responses against *Ng* (22, 46, 65–67, 69). Accordingly, we incorporated CpG ODN 2395, a TLR9 agonist that promotes Th1 polarization, into the BamA mRNA formulation for the IN immunization/challenge study.

**Figure 3.**
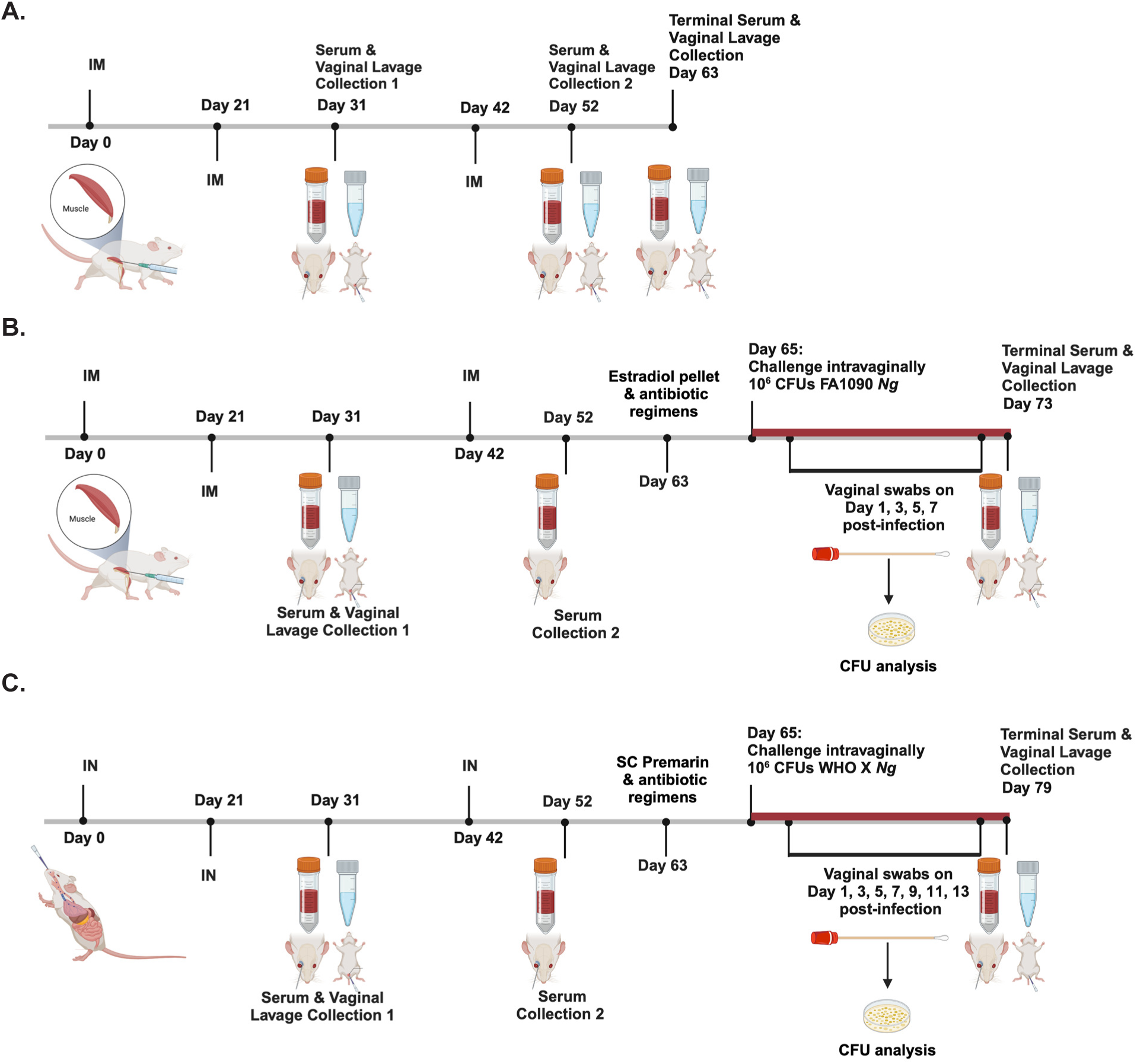
Experimental design of the pilot immunogenicity, IM immunization/challenge, and IN immunization/challenge studies. **A**, Pilot intramuscular (IM) immunogenicity study. Female BALB/c mice (*n*=6-7 per group) received IM immunizations with BamA mRNA-LNP (10 µg), EGFP mRNA-LNP (10 µg), or PBS on days 0, 21, and 42. Sera and vaginal lavages were collected on days 31, 52, and 63. **B**, IM immunization/challenge study. Mice (*n*=18-22/group) received IM immunizations on days 0, 21, and 42. Sera were collected on days 31, 52, and 73 and vaginal lavages on days 31 and 73. Mice received a 17-β estradiol pellet and an antibiotic regimen prior to an intravaginal challenge with 10^6^ CFU of *N. gonorrhoeae* FA1090 on day 65. Vaginal swabs collected on days 1, 3, 5, and 7 post-infection were plated on antibiotic-supplemented GC agar for CFU enumeration. **C**, IN immunization/challenge study. Mice (*n*=15-25/group) received intranasal (IN) immunizations of BamA mRNA-LNP (10 µg), BamA mRNA-LNP (10 µg) + CpG (15 µg), CpG (15 µg), or PBS on days 0, 21, and 42. Sera were collected on days 31, 52, and 79 and vaginal lavages on days 31 and 79. Mice received subcutaneous Premarin and an antibiotic regimen prior to intravaginal challenge with 10^6^ CFU *N. gonorrhoeae* WHO X on day 65. Vaginal swabs collected on days 1, 3, 5, 7, 9, 11, and 13 post-infection were plated for CFU enumeration.

In the pilot study, mouse cohorts (*n*=6-7) were immunized with three doses of PBS, EGFP mRNA LNPs, or BamA mRNA LNPs (**Fig. 3A**), and serum and vaginal lavage samples were collected on days 31, 52, and 63. In the follow-up, independent IM and IN immunization and challenge studies, mice (*n*=15-25/cohort) received PBS, EGFP mRNA LNPs, or BamA mRNA LNPs, or PBS, CpG LNPs, BamA mRNA LNPs, or BamA mRNA +CpG LNPs on days 0, 21, and 42 (**Fig. 3B and C**, respectively). Sera were collected on days 31 and 52, and at study termination on day 73 (for IM immunization) and on day 79 (for IN vaccine administration), while vaginal lavages were collected on days 31 and 73/79. In the IM immunization/challenge, we monitored *Ng* FA1090 infection for 7 days, whereas in the IN immunization/challenge study, we extended monitoring of *Ng* WHO X infection to 13 days (**Fig. 3C**).

### Immunization with BamA mRNA-LNPs elicits systemic and mucosal antigen-specific antibodies in mice

To evaluate whether different vaccine formulations delivered IM or IN induced antigen-specific systemic and mucosal IgG and IgA antibodies, immunoblotting experiments were conducted using pooled mouse sera or vaginal lavages (**Fig. 4**). In both studies with IM vaccine administration, serum IgG and IgA cross-reacted with *Ng* BamA expressed in *E. coli* (*E. coli* pBamA) and in whole cell lysates of isogenic *Ng* FA1090, as well as the *Ng* 2016 WHO reference strains (**Fig. 4A**). In contrast, the IN-administration route failed to elicit BamA-specific serum IgA regardless of whether mice were immunized with BamA mRNA LNPs or BamA mRNA+CpG LNPs. The IM route also elicited vaginal BamA-specific IgG and IgA, whereas no signal was detected in pooled vaginal lavages from mice immunized via the IN route (**Fig. 4B**). As expected, sera or vaginal lavages from control groups that received PBS, EGFP mRNA LNPs, or CpG LNPs did not show any BamA-specific IgG or IgA antibodies (**Fig. 4**).

**Figure 4.**
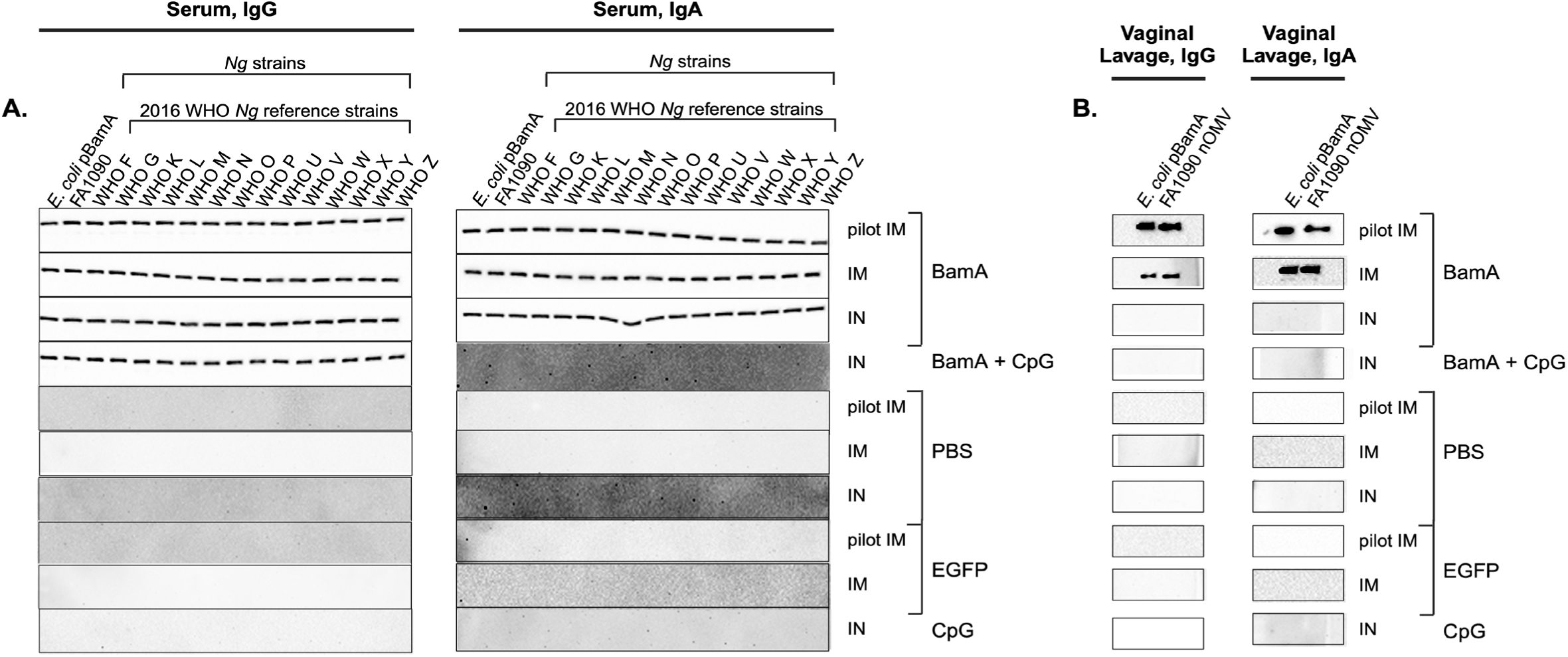
BamA mRNA-LNP immunization elicits systemic and mucosal antibodies that cross-react with diverse *N. gonorrhoeae* isolates. Pooled sera (**A**) and vaginal lavages (**B**) from mice immunized IM (pilot and IM challenge cohorts) or IN with BamA mRNA-LNP, BamA + CpG mRNA-LNP, EGFP mRNA-LNP, CpG, or PBS were analyzed by immunoblotting. **A**, Serum IgG (left) and IgA (right) on day 52 post-immunization, probed against recombinant BamA (*E. coli* pBamA), *N. gonorrhoeae* FA1090, and the 2016 WHO reference panel (WHO F, G, K, L, M, N, O, P, U, V, W, X, Y, Z). **B**, Vaginal IgG (left) and IgA (right) on day 63 (pilot IM cohort) or day 31 (IM challenge and IN cohorts), probed against *E. coli* pBamA and *N. gonorrhoeae* FA1090 native outer membrane vesicles (nOMVs). Rows are labeled by immunization route (pilot IM, IM, IN) and administered formulation.

### ELISA profiling of immunoglobulin classes induced by BamA mRNA LNPs immunization

ELISA analyses of the immunoglobulin classes revealed that in the pilot study, mice IM administered BamA mRNA LNPs had consistently significantly higher endpoint titers of serum IgG, IgG1, IgG2a, and IgA on day 63 compared with those that received EGFP mRNA LNPs or PBS (**Fig. 5**). For IgG, IgG1, and IgG3, two immunizations were sufficient to elicit high titers (**Fig. 5A**, **5B**, **5E**). On day 63, the BamA mRNA LNPs induced higher total IgG, geometric mean of 4.5×10^5^, compared to 7.4×10^2^ and 2.1×10^3^ for mice immunized with EGFP mRNA and PBS, respectively (**Fig. 5A**). The calculated geometric means for serum IgG1 and IgG2a were 9.4×10^4^ and 3.1×10^4^ by day 63 was 3.1×10^4^ for mice immunized with BamA mRNA LNPs compared to 1.0×10^1^, 1.0×10^1^, and 1.0×10^1^ and 2.9×10^1^, respectively, for mice that received EGFP mRNA LNPs and PBS (**Fig. 5B**, **C**). In the terminal sera, the IgG2a/IgG1 ratio in the vaccine group was 0.33, suggesting a Th2-biased immune response (**Fig. 5D**, (70, 71). Systemic IgA in the vaccinated animals reached significance by day 63 (**Fig. 5F**). Vaginal IgG was significantly increased in mice that received BamA mRNA LNPs on days 31 and 63, whereas IgA was significantly higher in these mice on day 52, in comparison to control groups (**Figs. 5G** and **H**, respectively).

**Figure 5.**
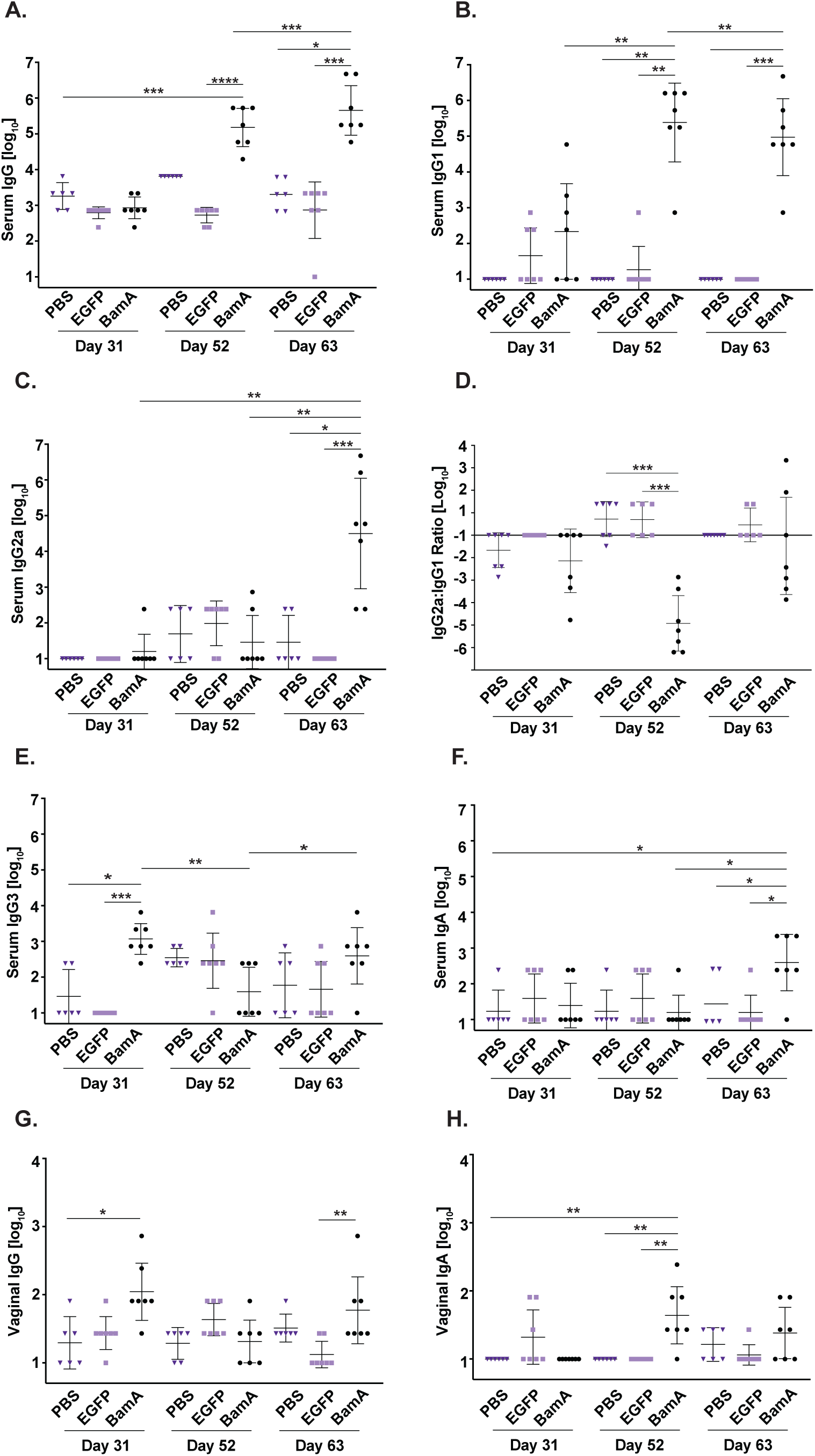
IM BamA mRNA-LNP immunization elicits sustained BamA-specific systemic and mucosal antibody responses in the pilot study. Sera and vaginal lavages from BALB/c female mice (*n*=6-7/group) immunized IM with BamA mRNA-LNP, EGFP mRNA-LNP, or PBS were assayed by ELISA against *N. gonorrh*oeae FA1090 nOMVs on days 31, 52, and 63 post-immunization. Serum IgG (**A**), IgG1 (**B**), IgG2a (**C**), IgG2a:IgG1 ratio, calculated as log_10_(IgG2a) − log10(IgG1) (**D**), IgG3 (**E**), and IgA (**F**); vaginal IgG (**G**) and IgA (**H**). Endpoint titers are shown as geometric means with 95% confidence intervals. Comparisons were performed using the Kruskal–Wallis test with Dunn’s multiple comparisons post hoc test. \**p* < 0.05, \*\**p* < 0.01, \*\*\**p* < 0.001, \*\*\*\**p* < 0.0001.

Subsequently, we analyzed antibody responses elicited by IM and IN vaccination in the two challenge studies (**Fig. 6**). Overall, three IM doses of BamA mRNA LNPs were sufficient to elicit significantly higher systemic IgG, IgG1, IgG2a, and IgG3 compared to the respective antibody titers in mice that received PBS or EGFP LNPs (**Figs. 6A**-**D**). The geometric mean of IgG1 increased by ∼700 (696.6-fold) on day 52 (after the second boost) compared to day 31 (**Fig. 6B**). There was also induction of serum IgA levels, which were 5-15-fold higher than those in control groups (**Fig. 6E**). IM BamA LNPs induced mild Th1 at d31 (GM = 1.28) and strong Th2 post-boost (GM = 0.011) as suggested by the IgG1/IgG2a ratios (**Fig. S2A**). Vaginal IgG and IgA levels were not significantly different from those in the control groups on day 31, consistent with the data from the pilot experiment (**Fig. S2B** and **C**).

**Figure 6.**
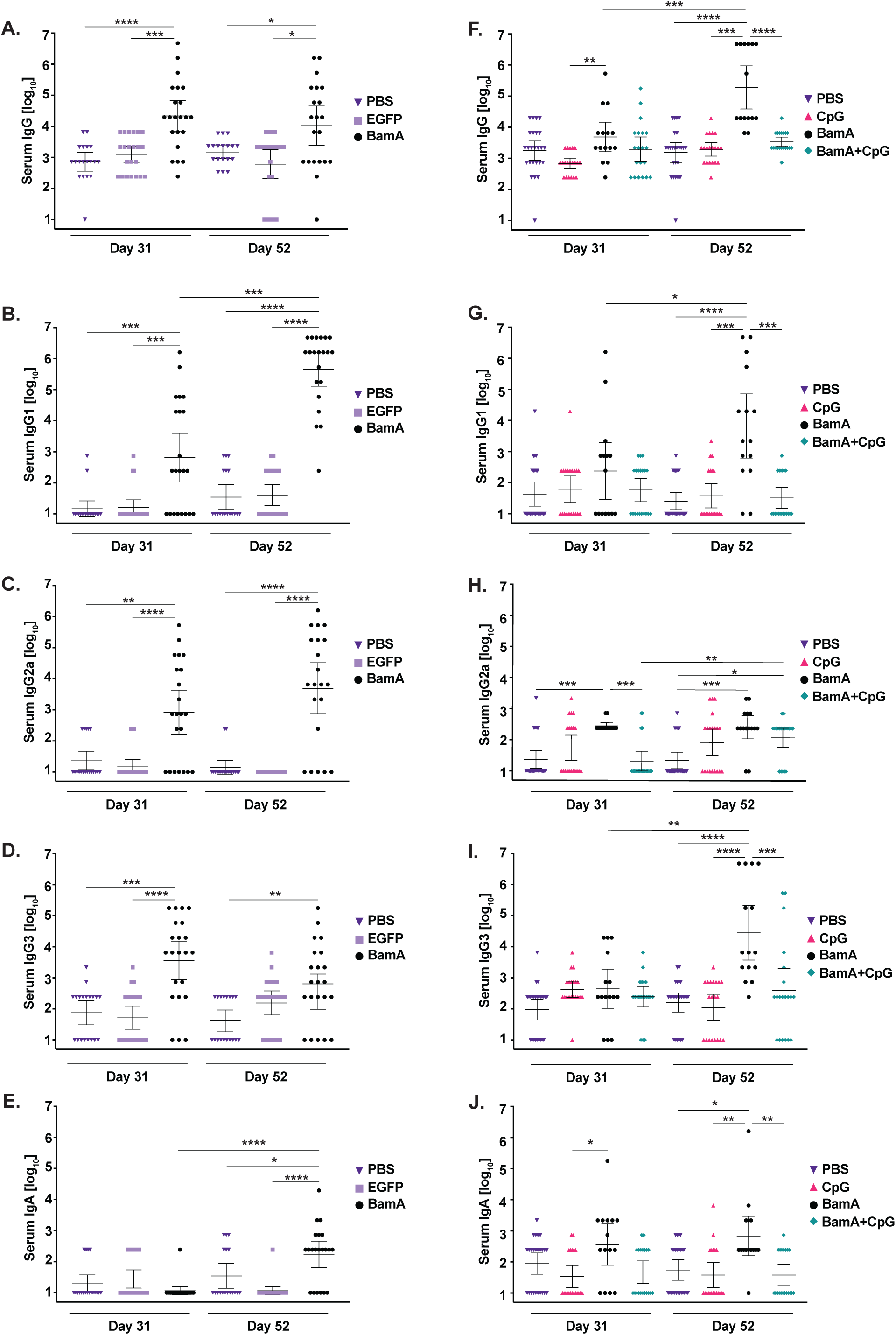
IM and IN BamA mRNA-LNP immunization elicit robust BamA-specific serum antibody responses in the immunization/challenge studies. Sera from mice immunized IM (**A**–**E**) with PBS, EGFP mRNA-LNP, or BamA mRNA-LNP or IN (**F**–**J**) with PBS, CpG, BamA mRNA-LNP, or BamA + CpG mRNA-LNP were assayed by ELISA against *N. gonorrhoeae* FA1090 nOMVs on days 31 and 52 post-immunization. Serum IgG (**A**, **F**), IgG1 (**B**, **G**), IgG2a (**C**, **H**), IgG3 (**D**, **I**), and IgA (**E**, **J**). Endpoint titers are shown as geometric means with 95% confidence intervals. Comparisons were performed using the Kruskal–Wallis test with Dunn’s multiple comparisons post hoc test. \**p* < 0.05, \*\**p* < 0.01, \*\*\**p* < 0.001, \*\*\*\**p* < 0.0001.

Two IN immunizations with the BamA mRNA LNPs vaccine elicited significantly higher levels of total serum IgG, IgG2a, and IgA compared to the BamA mRNA+CpG LNPs, CpG LNPs, and PBS, whereas the third immunization led to a further increase in IgG, IgG1, and IgG3 (**Fig. 6A-C**). The third IN dose boosted systemic IgG in mice vaccinated with BamA mRNA LNPs, but not in those administered the same vaccine adjuvanted with CpG (**Fig. 6F**), and the geometric mean titers were 1.9×10^5^, which was significant compared to 3.39×10^3^, 1.96×10^3^, and 1.55×10^3^ in mice that were administered BamA mRNA+CpG LNPs, CpG LNPs, or PBS, respectively. The serum IgG1 responses were 206-, 174-, and 260-fold higher than those of mice immunized with BamA mRNA+CpG LNPs, CpG LNPs, and PBS, respectively (**Fig. 6G**). The GM titer of serum IgG2a in BamA mRNA LNPs-immunized mice was 2.84×10^2^, which was significantly increased compared to that in mice immunized with BamA mRNA+CpG LNPs (2.11×10¹) and PBS (2.35×10¹). However, the difference was not statistically significant when compared to mice immunized with CpG alone (5.53×10^1^; **Fig. 6H**). Mice immunized with BamA mRNA+CpG LNPs had increased IgG2a antibody levels on day 52 compared to day 31; however, this increase was not significant relative to those in animals that received BamA mRNA LNPs or CpG LNPs. IN-delivered BamA LNPs showed a Th1 bias at d31 (GM = 1.92), which reversed to a strong Th2 bias post-boost at d52 (GM = 0.040). The BamA+CpG LNPs IN group was Th2 at d31 but reversed to the Th1 response post-boost (GM = 3.79). Further, sera obtained from mice immunized with BamA mRNA LNPs showed a significant increase in IgG3 with a geometric mean of 2.84×10^4^, compared to 3.89×10^2^,1.11×10^2^, and 1.61×10^2^ in animals immunized with BamA mRNA+CpG LNPs, CpG LNPs, and PBS, respectively (**Fig. 6I**). Systemic IgA in mice that received IN BamA mRNA LNPs were 18-, 17-, and 12-fold higher compared to IgA levels in mice administered with BamA+CpG LNPs, CpG LNPs, and PBS, respectively (**Fig. 6J**). In both vaccine groups, the IN-vaccine administration failed to induce vaginal IgG and IgA on day 31 (**Fig. S2E**-**F**).

### Intramuscular or intranasal immunization with BamA mRNA does not accelerate *Ng* clearance in mice

We assessed protection after IM and IN immunization against challenge with *Ng* FA1090 and a ceftriaxone-resistant, *Ng* WHO X (72, 73), respectively. Neither route nor vaccine formulation accelerated clearance as demonstrated on the Kaplan-Meier curves (**Fig. 7**). By day 7 in the IM study, 52.32%, 40.17%, and 62.82% of mice remained infected in the BamA mRNA, EGFP mRNA, and PBS groups, respectively (**Fig. 7A**), and three IN doses of BamA mRNA or BamA+CpG resulted in no statistically significant clearance of *Ng* WHO X on any of the thirteen days compared with controls (**Fig. 7D**). Bioburden was likewise unaffected by either route. Intramuscular immunization did not reduce CFU/mL (log_10_) BamA mRNA, 3.8 ± 1.72; EGFP mRNA, 4.11 ± 1.62; PBS, 3.55 ± 1.67; **Fig. 7B**), and the corresponding area under the curve (AUC) of cumulative CFU/mL (log_10_) was not diminished in vaccinated mice (**Fig. 7C**). Intranasal immunization similarly yielded no significant reduction in bioburden or AUC (**Fig. 7E**), with AUC of log_10_ CFU/mL of 12.6 ± 4.07, 16.16 ± 1.54, 12.14 ± 1.91, and 9.72 ± 2.97 for the BamA+CpG, BamA mRNA, CpG, and PBS groups, respectively (**Fig. 7F**).

**Figure 7.**
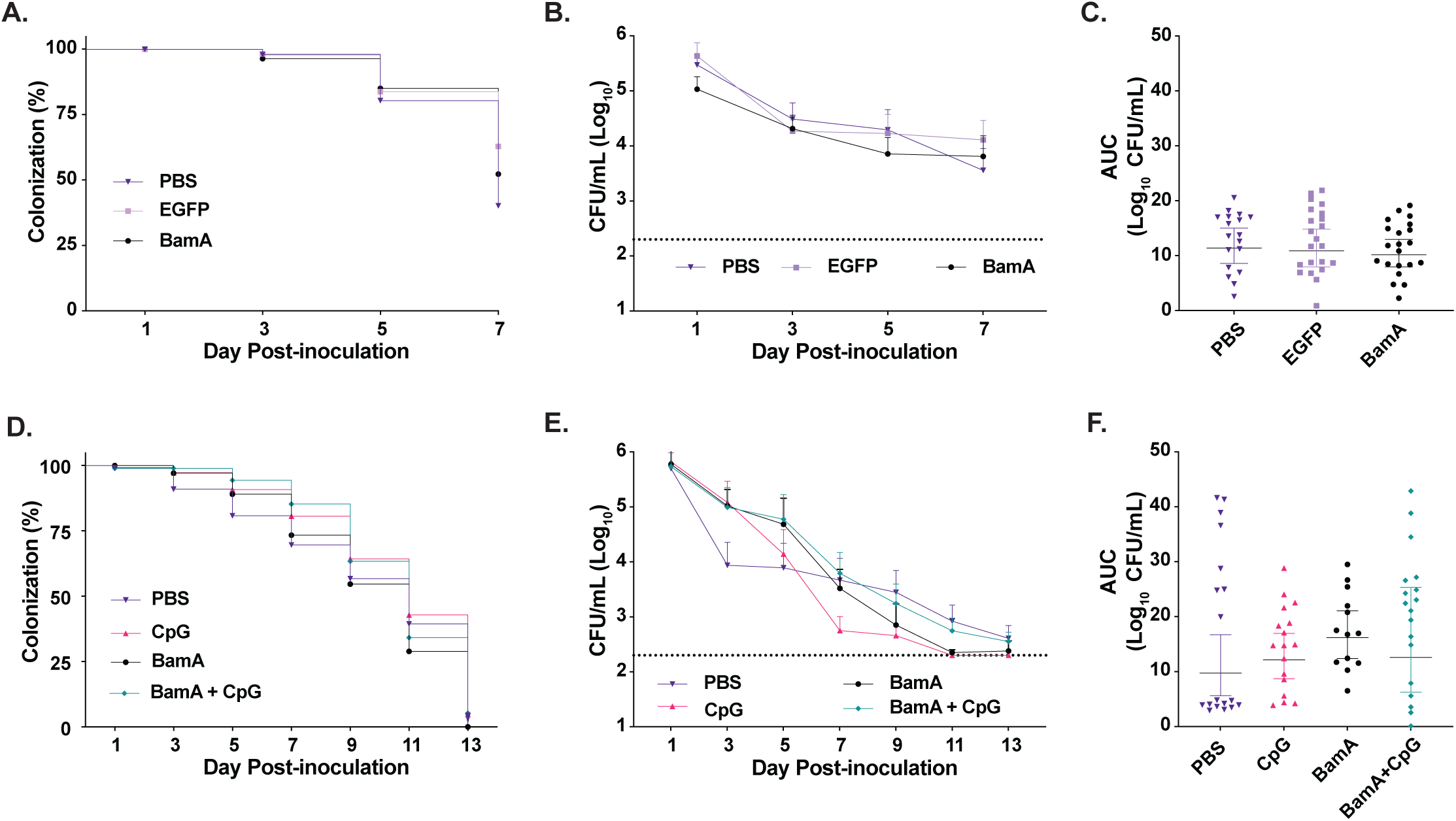
IM or IN BamA mRNA-LNP immunization does not accelerate *N. gonorrhoeae* clearance or reduce bioburden in the female mouse lower genital tract infection model. Female BALB/c mice (*n*=18-22/group) were immunized IM (**A**–**C**) with BamA mRNA-LNP, EGFP mRNA-LNP, or PBS and challenged intravaginally with 10^6^ CFU *N. gonorrhoeae* FA1090, or immunized IN (*n*=15-25/group; **D**–**F**) with BamA mRNA- LNP, BamA+CpG mRNA-LNP, CpG LNP, or PBS and challenged with 10^6^ CFU *N. gonorrhoeae* WHO X. Percentage of mice colonized over time (**A**, **D**), bacterial burden in individual mice (log_10_ CFU/mL) at each collection day (**B**, **E**), and cumulative bacterial burden (AUC, log_10_ CFU/mL) (**C**, **F**). Dotted lines in B and E mark the limit of detection (200 CFU/mL). Survival curves were compared using the log-rank (Mantel–Cox) test, bacterial burden at each time point by two-way ANOVA with Bonferroni correction, and AUC by the Kruskal–Wallis test with Dunn’s multiple comparisons. Bars show median with interquartile range.

To monitor polymorphonuclear leukocyte (PMN) influx, we scored the percentage of PMN per 100 cells in vaginal swabs (51). AUC PMN did not differ across groups by either route (intramuscular: 9.21 ± 1.38%, 9.25 ± 1.48%, and 9.68 ± 1.34% for BamA mRNA, EGFP mRNA, and PBS, **Fig. S3A**; intranasal: 15.39 ± 1.26%, 15.68 ± 1.18%, 14.8 ± 1.22%, and 14.99 ± 1.34% for BamA+CpG, BamA mRNA, CpG, and PBS over the thirteen days following challenge, **Fig. S3B**).

Together, these studies demonstrated that although both IM and IN BamA mRNA vaccines induced BamA-specific antibody responses, and IM delivery engaged a mucosal route, neither regimen accelerated clearance, reduced bioburden, or diminished PMN influx relative to PBS-, EGFP mRNA-, or CpG-immunized mice.

### BamA mRNA-LNP immunization fails to induce serum bactericidal activity

Finally, we determined the human complement-dependent serum bactericidal activity (hSBA) elicited by mRNA-LNP vaccination using pooled murine sera and IgG/IgM-depleted normal human serum (NHS) as the complement source. In the IM study, pooled sera from the PBS, EGFP mRNA, and BamA mRNA groups yielded SBA titers of 2048, 1024, and 2048, respectively (**Table 4**). In the intranasal study, pooled sera from the PBS, CpG-LNP, BamA mRNA-LNP, and BamA mRNA+CpG-LNP groups yielded SBA titers of 4096, 2048, 2048, and 2048, respectively (**Table 4**). Bactericidal titers in all vaccinated groups were comparable to or lower than those of the unimmunized PBS control, indicating that neither the vaccine formulation nor the immunization route elicited hSBA activity above background.

**Table 4.**
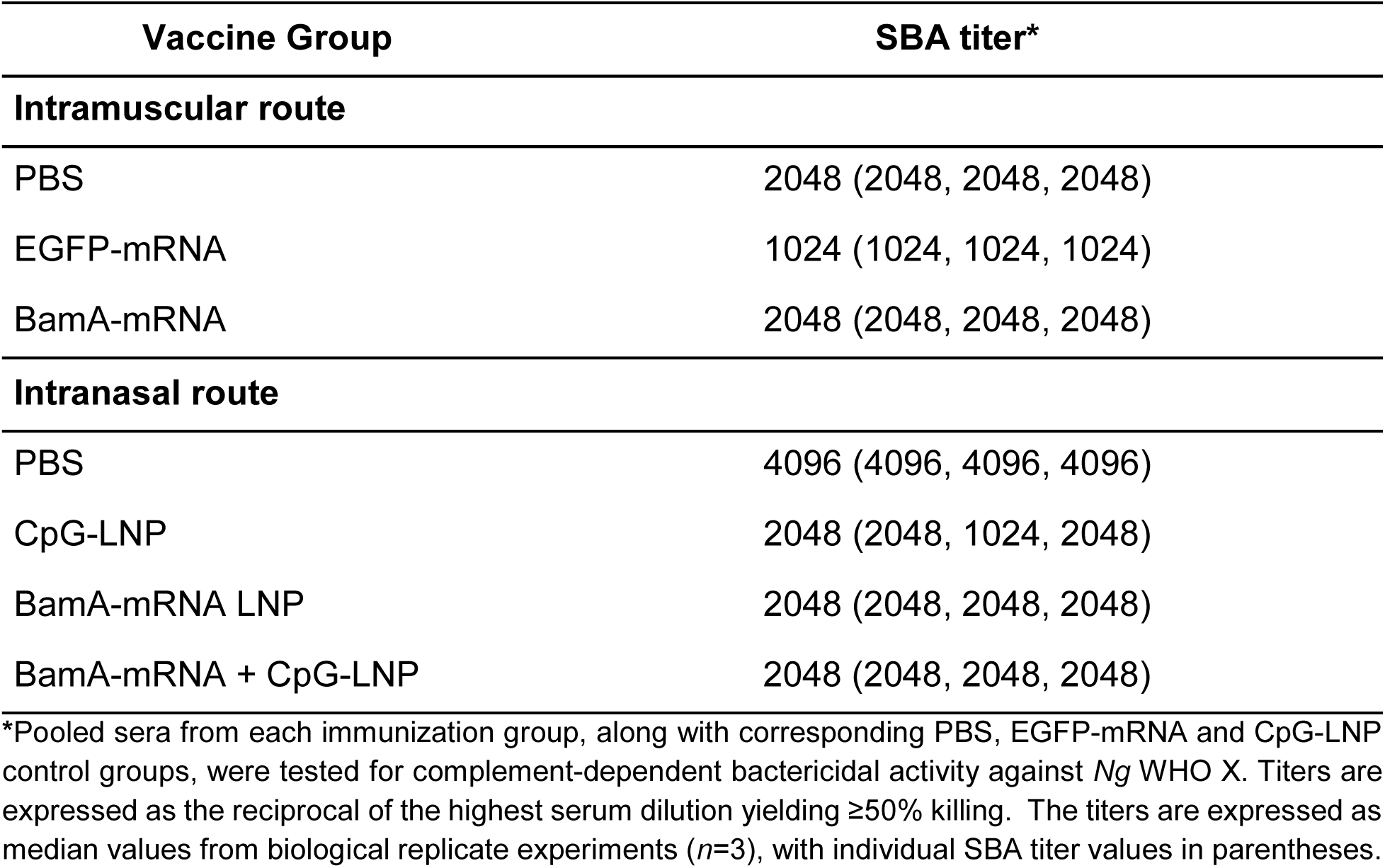
Complement-dependent serum bactericidal activity of pooled murine sera from mice immunized with BamA mRNA-LNP via intramuscular and intranasal routes against *N. gonorrhoeae* WHO X.

## Discussion

The GoGoVax double-blind, randomized controlled trial in a high-risk Australian population found no protective efficacy of 4CMenB against gonorrhea (74–76). Concurrently, the only gonococcal-specific OMV vaccine candidate to reach clinical evaluation (GMMA; GSK) was discontinued before efficacy data were published, leaving no late-stage, dedicated gonorrhea vaccine in the pipeline (77, 78). Together, these recent and historical vaccine setbacks (22) highlight the challenges of developing vaccines against *Ng* and underscore the critical need for purpose-built gonococcal vaccines capable of eliciting protective immunity.

The mRNA-lipid nanoparticle (LNP) platform has transformed vaccine development against viral diseases and is advancing rapidly into bacterial pathogens, with more than a dozen preclinical studies and two clinical candidates reported to date against *B. burgdorferi* (NCT05975099) and *M. tuberculosis* (NCT05537038, NCT05547464) (5). Here, we present the first evaluation of an mRNA-LNP vaccine against *Ng* that encodes the highly conserved outer membrane antigen BamA. Three independent preclinical studies, a pilot IM immunization and two immunization/challenge experiments using IM and IN routes, established that BamA mRNA-LNPs were strongly immunogenic and elicited cross-reactive antibodies against geographically and genetically diverse *Ng* isolates. Three IM doses generated robust serum IgG, IgG1, IgG2a, and IgA, together with mucosal vaginal IgG and IgA detectable by both immunoblotting and ELISA, whereas IN administration produced significant systemic IgG, IgG1, IgG2a, IgG3, and IgA responses. These antibody responses did not translate into accelerated bacterial clearance, reduced bioburden, or hSBA activity above the baseline of control sera. Two convergent, mRNA-platform-specific factors provide a potential explanation for the dissociation between BamA-specific antibody induction and protection. First, our *in-silico* analysis revealed a B-cell epitope distribution consistent with the immunogenic but non-bactericidal phenotype, with the highest-scoring linear and discontinuous epitopes localized to the periplasmic POTRA domains and the membrane- embedded β-barrel rather than to the surface-exposed extracellular loops L1, L5, and L8 (**Fig. 1**). Antibodies directed at POTRA and β-barrel regions recognize denatured or lysed bacteria on immunoblots but engage few accessible targets on the intact gonococcal surface. Second, and central to mRNA vaccine design, bacterial antigens expressed in mammalian cells undergo host post-translational modifications (PTMs), particularly N- and O-linked glycosylation, which can alter or mask immunogenic epitopes (5). The mIgκ signal peptide we incorporated routed BamA through the ER/Golgi secretory pathway, enabling efficient secretion into the supernatants of HEK293 cells (**Fig. S1**) and likely positioning the protein for glycosylation. Consistent with this, mRNA-encoded BamA carrying mIgκSP showed a slight upward shift in molecular mass relative to the SP-devoid construct, indicative of mammalian processing. We incorporated the murine Igκ as it enhanced antigen expression and processing in host cells and increased vaccine efficacy (57–60). However, two other precedents bear contrasting results. An mRNA vaccine encoding *Y. pestis* F1 fused to a mammalian Igκ signal peptide elicited cellular but no humoral responses, attributed to PTMs that masked critical immunogenic epitopes; protection was achieved when F1 was expressed without a signal peptide and reached the extracellular space via unconventional secretion (13). For *M. tuberculosis* Ag85A, host N-glycosylation similarly impaired humoral and cellular immunity to a nucleic acid-encoded antigen, and immunogenicity was restored by mutagenesis of the N- glycosylation site, a finding that has been hypothesized to contribute to the clinical failures of the viral-vectored MVA85A and AERAS-402 TB candidates (79). For BamA, mammalian PTMs may further obscure the already-limited set of surface-loop B-cell epitopes, compounding the accessibility problem.

The mRNA-LNP platform’s natural strength in eliciting T-cell responses offers a complementary, and arguably more tractable, route to BamA-mediated protection. Cytosolic translation of mRNA-encoded antigens favors proteasomal degradation and MHC-I presentation (5), and our prediction identified 28 high-affinity MHC-I epitopes, including peptides in the surface-exposed loops L1 (score 0.953), L5 (0.817), and L8 (0.981). The same loops that are inaccessible to serum antibody may therefore be processed and presented to CD8+ T cells on infected mucosal cells. This T cell-centric mode of protection has precedent in bacterial mRNA vaccines: an mRNA construct encoding the bacterial surface protein LMON_0149 conferred CD8+ T cell-dependent protection against *Listeria monocytogenes* even with limited humoral response and encoded antigen instability (80), and an LNP-mRNA tuberculosis vaccine drove polyfunctional Th1 CD4+ T cells in blood and lungs (81). Cellular immunity was not directly assessed in our study, and the Th2-skewed IgG2a/IgG1 ratios provide only an indirect readout. Incorporating intracellular cytokine staining, ELISpot, and assessment of tissue-resident memory T cells alongside functional antibody readouts will be a priority for next-generation candidates and will clarify whether T cell-mediated immunity can be leveraged for BamA-specific protection.

These findings define a clear, mRNA-specific path forward. To address the dual challenges of epitope accessibility and PTMs, engineering the BamA antigen is the primary priority. Bioinformatic prediction of N- and O-linked glycosylation sites with NetNglyc and NetOglyc, followed by site-directed mutagenesis to eliminate host glycosylation while preserving native folding, can restore native loop conformation and immunogenicity (79). Alternative antigen design, including soluble loop-only or membrane-anchored constructs that prominently display L1, L5, and L8 without the buried POTRA scaffold, could redirect the antibody response toward functionally accessible targets. Routing strategies offer another way for optimization. Removing the signal peptide to drive cytosolic expression (favoring MHC-I presentation), incorporating LAMP1 or MHC-I trafficking signals (enhancing both MHC-I and MHC-II presentation), or including an IgG-Fc fragment (extending plasma half-life and targeting MHC-II via Fcγ receptor-mediated endocytosis) have each produced antigen-specific gains in bacterial mRNA systems (5, 13, 17). Construct-level enhancements include codon optimization for mammalian translation, GC content adjustment, which improved longevity and translation in large-animal studies, and self-amplifying mRNA (SAM) backbones, which drove protection against group A and B Streptococcus and *Y. pestis* at lower doses (5, 10, 14, 18, 82–84). At the formulation level, the LNP itself provides opportunities to incorporate immunostimulatory glycolipids, for instance α-galactosyl ceramide (α-GalCer), which enhanced Th1 priming at mucosal surfaces and has demonstrated activity against *Helicobacter pylori* (85), and engineered LNP adjuvants are emerging as a strategy to broaden innate immune activation in mRNA vaccines (5). For mucosal protection, heterologous IM-prime/IN-boost schedules outperformed homologous IM regimens for SARS-CoV-2 mucosal immunity (86) and warrant evaluation. Finally, mRNA’s defining advantage, concurrent encoding of multiple antigens, supports multivalent BamA constructs that include additional conserved gonococcal antigens identified by our proteomics-driven reverse vaccinology (32, 33, 87, 88) to broaden epitope coverage and exploit immunodominance hierarchies, an approach that has produced cross-species protection in multivalent *Streptococcus* and *Clostridioides difficile* mRNA vaccines (15, 82).

This first evaluation of an mRNA vaccine against *Ng* establishes the platform as a new approach for gonococcal antigen testing. It demonstrates that mRNA-LNPs elicit robust, cross-reactive humoral responses against a highly conserved gonococcal antigen, suggests mammalian post-translational processing as a likely contributor to the immunogenic-but-non-protective phenotype, and defines testable modifications to the construct, adjuvant, and route. The speed, flexibility, and scalability that have made mRNA-LNPs the platform of choice for next-generation vaccines (5) are particularly well suited to the iterative antigen optimization required by gonococcal vaccine development. Building on the framework established here, refined BamA constructs and multivalent combinations represent a promising direction toward a purpose-built mRNA vaccine against gonorrhea.

## Author Contributions

Conceptualization, A.E.S.; methodology, N.W., N.T.V.M., J.K., A.C., R.K., R.A.Z., F.G.M. and G.S.; investigation, N.W., A.C., R.K. and F.G.M.; formal analysis, N.W. and A.C.; resources, G.S. and A.E.S.; writing—original draft preparation, N.W.; writing—review and editing, all authors; supervision, A.E.S. and G.S.; funding acquisition, A.E.S. All authors have read and agreed to the published version of the manuscript.

## Acknowledgments

We thank Lixin Li and Yujuan Song for technical assistance with animal experiments. The authors thank the staff of the Oregon State University Laboratory Animal Resources Center for animal husbandry support. Funding was provided to Dr. Aleksandra Sikora through grant R01-AI117235 from the National Institute of Allergy and Infectious Diseases, National Institutes of Health, and the Oregon State University Advantage Accelerator. The funders had no role in the design of the study; in the collection, analyses, or interpretation of data; in the writing of the manuscript; or in the decision to publish the results.

## Institutional Review Board Statement

The animal study protocol was approved by the Institutional Animal Care and Use Committee of Oregon State University (protocol #IACUC2021-0244). Animal experiments were performed in accordance with the Guide for the Care and Use of Laboratory Animals (NIH Publication No. 85-23) and the 2020 AVMA Guidelines for the Euthanasia of Animals.

## Informed Consent Statement

Not applicable.

## Data Availability Statement

The data presented in this study are available within the article and its Supplementary Materials. Additional data are available from the corresponding author upon reasonable request.

## Supplementary Figures

**Supplementary Figure S1. Cellular localization of mammalian-expressed BamA. A**, Schematic of pSecTag2A-BamA constructs without (left) and with (right) the mouse IgGκ signal peptide (SP). HEK293 cells were transfected with lipofectamine alone (mock), empty pSecTag2A, pSecTag2A-BamA, or pSecTag2A-IgGκ-BamA, and BamA expression in cell pellets (**B**) and pyrogallol red–molybdate–methanol (PRMM)- precipitated culture supernatants (**C**) was detected by immunoblotting with polyclonal rabbit anti-BamA antisera at two protein loadings (5 µg, 10 µg). BamA lacking the SP migrated at ∼88 kDa; the IgGκ-fused construct showed an upward molecular mass shift consistent with mammalian post-translational modification. Secreted BamA was detected only in supernatants of cells transfected with pSecTag2A-IgGκ-BamA.

**Supplementary Figure S2. IgG2a:IgG1 ratios and mucosal antibody responses in the IM and IN immunization/challenge cohorts.** Sera and vaginal lavages from mice (*n*=18-22/group) in the IM (A–C) and IN (*n*=15-25/group; **D**–**F**) immunization/challenge studies were assayed by ELISA against *N. gonorrhoeae* FA1090 nOMVs. IgG2a:IgG1 ratio (log_10_) on days 31 and 52 post-immunization (**A**, **D**); vaginal IgG on day 31 (**B**, **E**); vaginal IgA on day 31 (**C**, **F**). IM cohorts: PBS, EGFP mRNA-LNP, BamA mRNA-LNP. IN cohorts: PBS, CpG, BamA mRNA-LNP, BamA + CpG mRNA-LNP. Endpoint titers are shown as geometric means with 95% confidence intervals. Comparisons were performed using the Kruskal–Wallis test with Dunn’s multiple comparisons post hoc test. \**p* < 0.05, \*\*\**p* < 0.001.

**Supplementary Figure S3. Vaginal polymorphonuclear leukocyte (PMN) responses following bacterial challenge.** Vaginal swabs collected during the IM (A) and IN (B) immunization/challenge studies were stained with Wright’s stain, and PMNs were enumerated as a percentage of 100 total cells. Cumulative AUC values for % PMNs are shown across the 7-day (IM, *N. gonorrhoeae* FA1090) or 13-day (IN, *N. gonorrhoeae* WHO X) infection course. IM cohorts: PBS, EGFP mRNA-LNP, BamA mRNA-LNP (**A**). IN cohorts: PBS, CpG, BamA mRNA-LNP, BamA + CpG mRNA-LNP (**B**). Bars show median with interquartile range. No significant differences were detected by the Kruskal– Wallis test.

**Supplementary Figures S4-11. Uncropped and unprocessed immunoblot images supporting Figure 4**. Pooled sera and vaginal lavages from mice immunized IM (pilot and IM challenge cohorts) or IN with BamA mRNA-LNP, BamA + CpG mRNA-LNP, EGFP mRNA-LNP, CpG, or PBS were immunoblotted for BamA-specific IgG and IgA against recombinant BamA, *N. gonorrhoeae* FA1090, the 2016 WHO reference panel (serum), and FA1090 nOMVs (vaginal lavage). Molecular weight (kDa) is indicated to the left of each panel.

